# GOTFlow: Learning Directed Population Transitions from Cross-Sectional Biomedical Data with Optimal Transport

**DOI:** 10.64898/2026.03.16.712005

**Authors:** George Wright, Ethar Alzaid, Joanne Muter, Jan Brosens, Fayyaz Minhas

## Abstract

**Motivation:** Many biological and clinical processes are dynamic, yet most datasets are cross-sectional, capturing populations at discrete states rather than tracking individuals over time. This makes it difficult to quantify how populations change across developmental, physiological, or disease-associated conditions. Existing trajectory and transport-based methods often rely on fixed feature spaces, assumptions tailored to transcriptomic time-course data, or approximately linear progression, limiting their ability to model heterogeneous and unbalanced transitions across diverse biomedical modalities. Flexible methods are needed that can infer directed population-level change from cross-sectional data while retaining biological interpretability.

**Results:** We present GOTFlow, a framework for learning directed population transitions from cross-sectional biomedical data using graph-constrained optimal transport in a learned latent space. GOTFlow integrates representation learning with unbalanced optimal transport to jointly estimate embeddings and transport couplings between biological states. This enables hypothesis-driven modelling of progression structures while accommodating non-linear geometry, branching relationships, and changes in population mass. From the inferred transport plans, GOTFlow derives interpretable summaries of dynamics, including drift vectors quantifying transitions, and feature-level transported changes that highlight molecular drivers of progression. In synthetic data, GOTFlow recovered known transitions with strong agreement between inferred and ground-truth drifts. Across three biological applications, endometrial remodelling, breast cancer risk progression, and prion disease, GOTFlow identified state-to-state transitions and biologically meaningful feature shifts reflecting impaired decidualisation, increasing cancer risk, and neurodegenerative progression. These results establish GOTFlow as a general and interpretable framework for analysing directed population dynamics from cross-sectional data.

**Availability:** Code available at: https://github.com/wgrgwrght/GOTFlow

## Introduction

Biological and clinical systems are inherently dynamic. Patients progress through disease stages and tissue remodel in response to physiological cues, and molecular profiles shift as pathological states emerge or resolve. These transitions involve coordinated changes across multiple molecular and phenotypic layers, including gene expression, protein abundance, cellular composition, and tissue morphology. Although modern profiling technologies now capture such processes at scale (1; 10), most computational analyses still treat data as static collections of samples. Standard approaches such as clustering, dimensionality reduction, and differential testing primarily describe population structure and provide limited insight into how populations change across developmental, pathological, or clinical states.

Trajectory inference and pseudotime methods attempt to reconstruct biological progression from static observations by embedding samples in low-dimensional manifolds and ordering them along inferred paths (17). While useful for exploratory analysis, these approaches depend strongly on the chosen embedding and sampling density, and typically provide relative orderings of samples rather than explicit models of population-level transitions or quantitative measures of how populations move between states. Moreover, most trajectory inference methods were developed for single-cell transcriptomic datasets, where dense sampling across developmental or experimental conditions enables reconstruction of cell-state progressions.

RNA velocity estimates short-term transcriptional changes from spliced and unspliced transcript abundances to infer likely future cellular states (13). However, its reliance on transcriptomic kinetic information limits applicability across other modalities and many cohort-based biomedical studies. Optimal transport (OT) provides a broader alternative by estimating couplings between distributions and has been used to study biological population shifts and lineage relationships (18; 8; 5). Yet many OT-based methods operate in fixed feature spaces, such as PCA embeddings, where distances may not reflect biologically meaningful transition structure.

Many biomedical datasets are cross-sectional rather than longitudinal, meaning that samples represent heterogeneous populations observed under different biological or clinical states rather than repeated measurements of the same individuals. Modelling transitions between such populations therefore requires approaches that infer population-level dynamics without relying on explicit temporal tracking, motivating methods that operate on transitions between distributions of samples rather than trajectories of individual observations.

To address this gap, we introduce GOTFlow, a framework for modelling directed transitions between biological populations in high-dimensional data. GOTFlow combines representation learning with unbalanced OT to learn a latent geometry in which population-level flows between states are inferred along a user-defined directed state graph. This enables hypothesis-driven modelling of progression structures such as developmental stages, treatment responses, or disease risk strata while providing interpretable summaries of transition direction and feature-level change.

The key contributions of this paper are as follows:

1. **Graph-constrained representation learning with unbalanced optimal transport**. We introduce GOTFlow, a generalised OT framework for modelling transitions between heterogeneous biological populations from cross-sectional data. Unlike most OT trajectory methods designed for single-cell dynamics, GOTFlow supports cohort-level analyses where states correspond to biological or clinical conditions (e.g. timing, disease stages or treatment responses). A user-defined directed state graph encodes admissible transitions and enables modelling of branching, context-dependent, and unbalanced population changes.
2. **Interpretable summaries of population dynamics**. From inferred transport plans, GOTFlow derives barycentric drift vectors and feature-level transport summaries that quantify the direction and magnitude of population change, enabling comparative analysis of progression dynamics across biological or clinical conditions and highlighting molecular drivers associated with altered transitions.
3. **Validation across synthetic and biomedical datasets**. We evaluate GOTFlow on synthetic datasets with known transition structures and apply it to three biological studies: endometrial remodelling, breast cancer risk trajectories, and prion disease progression, demonstrating interpretable molecular shifts along population transitions.

## Methods

### Problem Formulation

GOTFlow models directional biological transitions, such as disease progression or tissue remodelling, as population-level flows between states in a learned embedding space. Rather than reconstructing trajectories of individual samples, the method seeks to explain how distributions of observations associated with different biological states relate to one another through transitions. For example, in cancer progression patients may be stratified into ordered risk states, and the goal is to characterise how molecular profiles evolve across this risk continuum.

Let 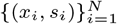 denote a dataset of *N* observations, where *x*_*i*_ ∈ 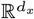 is a feature vector (e.g. gene expression profiles or histological image embeddings) and *s*_*i*_ ∈ *N* = {1, …, *K*} is a discrete state label corresponding to biological or clinical conditions such as time points, developmental stages, disease stages, or risk strata.

Admissible transitions between states are specified by a directed weighted graph with *L* edges between states 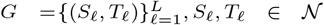 where each edge (*S, T*) represents a hypothesised transition between states (e.g. early*→*late or control*→*treatment). Edges may optionally be assigned weights *w*_*ST*_ ≥ 0 to control their contribution to the learning objective.

The goal of GOTFlow is to learn a representation *ϕ*_*θ*_ : 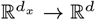 mapping each observation to a latent embedding *z*_*i*_ = *ϕ*_*θ*_(*x*_*i*_) such that the geometry of the latent space reflects the transition structure encoded by *G*. In particular, transitions permitted by the graph should correspond to efficient population flows between the associated states, whereas transitions not supported by the graph, or explicitly specified as incompatible, should incur higher cost. This encourages the learned representation to capture the progression structure encoded by *G* while discouraging spurious transitions (Fig. 1). As discussed below, this is achieved by a parameter encoding followed by unbalanced optimal transport in a single learning objective.

**Figure 1.**
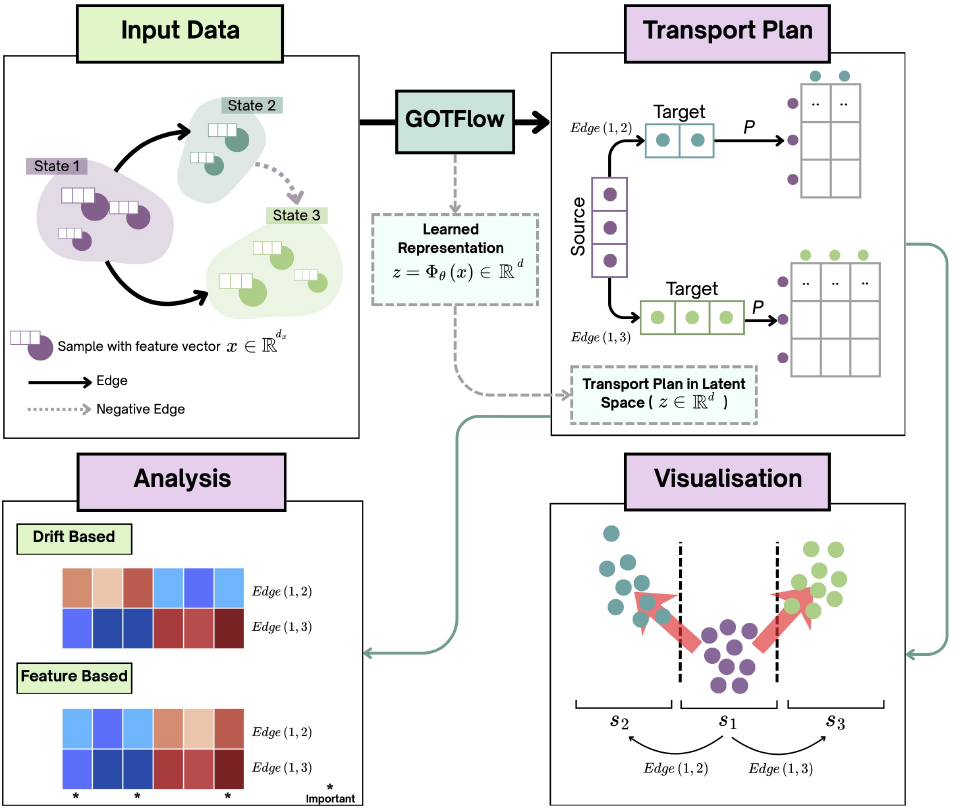
Overview of the GOTFlow framework. Samples are represented in their original feature space and assigned to discrete states connected by a user-defined directed graph specifying admissible transitions. GOTFlow jointly learns a representation and estimates optimal transport plans between connected states. The resulting transport plans are visualised and analysed to characterise state-to-state transitions, including drift vectors describing transition dynamics and feature-level attribution analyses identifying variables that change most strongly across edges.

### Latent Representation and Whitening

GOTFlow learns a parametric encoder *ϕ*_*θ*_ : 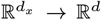 that maps each observation to a latent embedding *z*_*i*_ = *ϕ*_*θ*_(*x*_*i*_). The encoder’s parameters *θ* are trained jointly with the transport objective so that distances in the latent space reflect meaningful relationships between state populations and facilitate efficient transport along edges in the transition graph.

Directly optimising transport costs over unconstrained embeddings can lead to degenerate solutions, for example when the encoder collapses representations so that all samples become nearly identical and transport costs vanish. Moreover, optimal transport is sensitive to scale and anisotropy in the embedding. To stabilise optimisation and prevent such collapse, we apply global whitening to the latent representations before computing transport.

Let *µ*_*z*_ and Σ_*z*_ denote the mean and covariance of the embeddings {*z*_*i*_}. We define the whitening transform 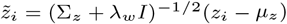, where *λ*_*w*_ > 0 is a small regulariser for numerical stability. Optimal transport computations are then performed in the whitened space 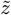. In practice, the whitening statistics are periodically recomputed from large minibatches and held fixed for several training epochs.

### Modelling Transitions with Unbalanced Optimal Transport

To model transitions between states, each state population is represented as a discrete empirical measure supported on the whitened latent embeddings. For a transition edge (*S, T*) ∈ *G*, let *I*_*S*_ = {*i* : *s*_*i*_ = *S*} and *I*_*T*_ = {*j* : *s*_*j*_ = *T*} denote the sets of samples belonging to the source and target states. The corresponding state populations are 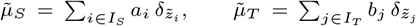, where *a*_*i*_, *b*_*j*_ ≥ 0 denote sample weights satisfying 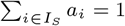 and 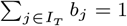.

Transitions between states are modelled as transport between these distributions. In biological systems, however, populations across states often differ in size or composition due to processes such as proliferation, cell death, or the emergence of new subpopulations. Enforcing strict mass conservation would therefore impose unrealistic one-to-one matching between samples. GOTFlow instead uses *unbalanced optimal transport* (UOT), which allows mass to be created or removed when moving from *S* to *T* and thus captures partial correspondence between populations.

For discrete measures supported on the embeddings 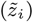, a transport plan is represented by a matrix 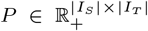 whose entries *P*_*ij*_ indicate how much mass moves from source sample *i* to target sample *j*. Given a coupling *P*, the unbalanced entropic OT functional is

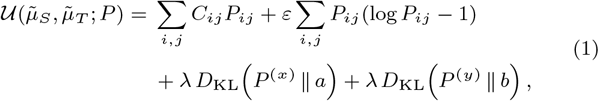

where 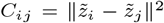 is the transport cost, *P* ^(*x*)^ = Σ*P*_*ij*_ and *P* ^(*y*)^ = Σ*P*_*ij*_ denote the row and column sums of *P*, and *D*_KL_ is the Kullback–Leibler divergence. The first term minimises the transport cost under the latent geometry, the entropic blur term (controlled by *ε*) promotes smooth and numerically stable couplings, and the KL penalties (weighted by *λ*) relax strict mass conservation by allowing deviations between transported and original masses.

To remove the entropic self-similarity bias, we define a Sinkhorn-divergence-style transition energy for edge (*S, T*) (7):

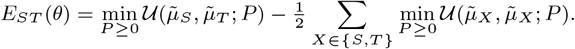

which measures the compatibility between the learned representation *ϕ*_*θ*_ and the given transition. The inner optimisation over *P* is performed using the Sinkhorn algorithm implemented in the GeomLoss library (7), yielding a differentiable estimate of *E*_*ST*_ (*θ*) for gradient-based optimisation of the latent embedding across all transitions.

### Final Training Objective of GOTFlow

The encoder parameters *θ* are learned so that the latent geometry makes given graph transitions transport-efficient while discouraging implausible ones. Intuitively, if state *S* represents an early biological condition and *T* a later stage, the embedding should place their populations so that the transport cost *E*_*ST*_ (*θ*) is small, while transitions to unrelated states incur higher cost. This optimisation has a bilevel structure: an inner optimal transport problem computes the transition energies *E*_*ST*_ (*θ*), while the outer optimisation updates the encoder parameters *θ* to favour valid transitions. This is achieved using two complementary objectives. First, OT energy minimisation encourages embeddings that reduce the transition energy *E*_*ST*_ (*θ*) along the edges (*S, T*) ∈ *G*. Second, a contrastive objective ensures that true transitions have lower energy than alternative destinations.

For a positive transition (*S, T*) ∈ *G*, we compare the OT energy of the true transition against a set of negative candidates. These negatives may either be transitions explicitly specified by the user as implausible or randomly sampled state pairs (*S, U*) for which no edge exists in the graph. Lower OT energy indicates a more plausible transition.

The resulting sampled softmax (InfoNCE) loss for edge (*S, T*) is

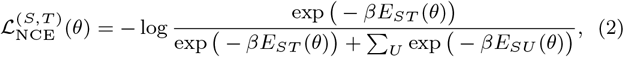

where *β >* 0 is an inverse temperature parameter controlling the sharpness of the comparison. The complete training objective aggregates these terms across all edges:

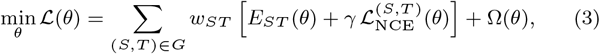

where *γ ≥* 0 controls the strength of the contrastive term and Ω(*θ*) denotes *𝓁*_2_ weight decay or sparsity regularisation.

This formulation subsumes several useful regimes. With fixed embeddings, optimisation reduces to estimating transport plans between state populations. When *θ* is learned jointly with transport, the embedding adapts to make valid transitions geometrically efficient. Depending on the mass regularisation parameter, the transport can operate in balanced or unbalanced regimes, enabling modelling of both conserved and non-conserved population dynamics. The contrastive term can be omitted (*γ* = 0) for purely transport-based alignment or included to enforce discriminative transition structure. Because the OT energies are computed using differentiable Sinkhorn iterations, the entire objective remains differentiable and can be optimised efficiently using GPU-based gradient methods.

### Drift Summaries and Feature-Level Interpretation

After training, GOTFlow uses the learned transport plans to summarise transition dynamics between states. For each directed edge (*S, T*) ∈ *G*, the optimal transport plan *P* defines how mass from samples in state *S* moves toward samples in state *T*. Each source sample *i* ∈ *I*_*S*_ is therefore associated with a barycentric projection onto the target population, 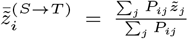 which induces a latent drift vector 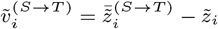.

When multiple transitions originate from state *S*, these edge-specific drifts are aggregated into a net drift summary

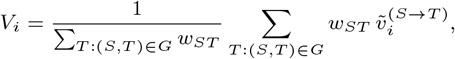

whose magnitude *∥V*_*i*_*∥* quantifies the overall transition intensity for sample *i*, highlighting individuals whose molecular profiles shift unusually strongly or weakly along a disease progression trajectory.

To interpret transitions in the original feature space, the same transport plan can be applied to the input features. For a source sample *i* on edge (*S, T*),

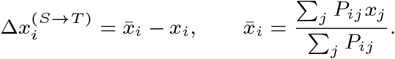

Aggregating 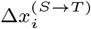 across source samples yields a transported-mass-weighted estimate of the average feature-level change associated with transition (*S, T*), highlighting molecular or phenotypic drivers of population shifts.

### Experimental Results and Applications

We evaluated GOTFlow on a synthetic benchmark and three biological datasets spanning controlled trajectories, endometrial remodelling, breast cancer risk progression, and prion disease. Across these settings, we tested whether GOTFlow can recover directed transition structure, quantify transition magnitude, and identify feature-level shifts associated with progression.

#### Synthetic Data Experiments

To validate whether GOTFlow can recover known transition structure, we generated a synthetic dataset of 100 samples in a 64-dimensional feature space with controlled perturbations producing known state-to-state displacements (see Supplementary Results). The data followed a bifurcating trajectory that split into two branches. Using the directed transition graph, GOTFlow recovered the expected transport structure, correctly capturing both divergence and merging transitions (Fig. 2). Per-edge OT feature-drift heatmaps (Supplementary Fig. S1) showed coherent signed feature shifts along the main trajectory and distinct drift signatures along each branch. Quantitative evaluation was performed using an independent held-out test set generated from the same process. The inferred drift vectors aligned closely with the ground-truth displacements, achieving a mean cosine similarity of 0.720 *±* 0.152. Additional ablation results and comparisons with classical OT showing advantages of key components of the pipeline are reported in Supplementary Table S1.

**Figure 2.**
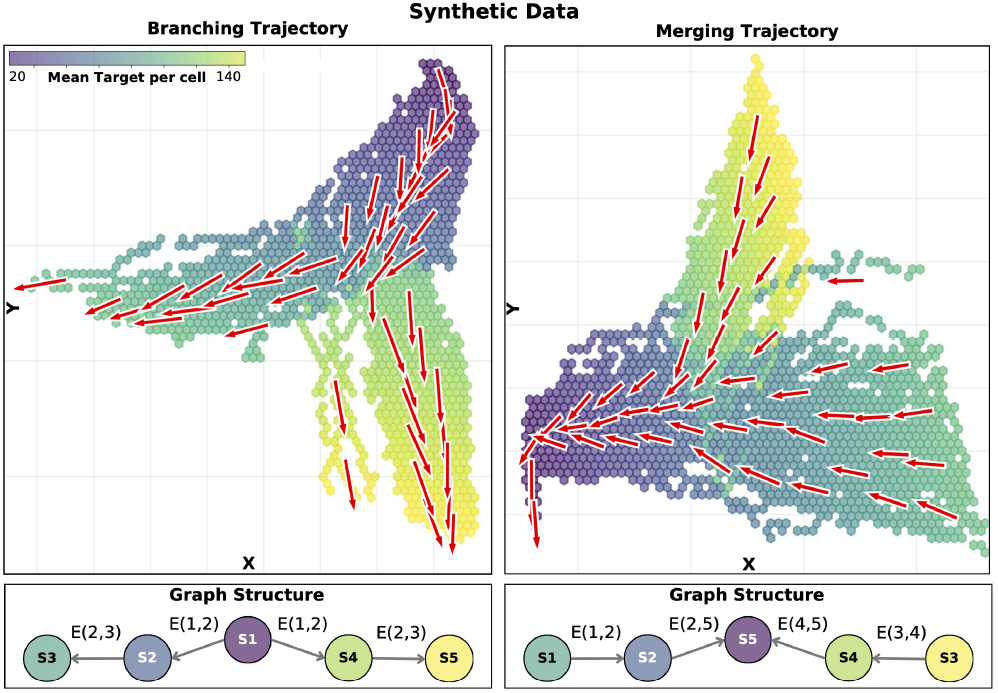
GOTFlow transport visualisation for the synthetic dataset. Samples are embedded in 2D and coloured by mean target per cell, red arrows indicate the inferred optimal-transport flow. Left: branching trajectory; right: merging trajectory. Each panel shows the corresponding state-transition graph, illustrating the states and directed edges used in the experiment.

#### Modelling Endometrial Gene Expression in the Luteal Phase

Following ovulation, the human endometrium undergoes coordinated transcriptional remodelling that prepares the tissue for embryo implantation. Disruption of these molecular transitions has been linked to implantation failure and recurrent pregnancy loss. We analysed gene expression profiles from 664 endometrial biopsies (514 miscarriage cases, 150 controls) obtained at the Implantation Research Clinic run by the Brosens lab. Samples were collected 4–12 days after the LH surge and grouped into early (LH+4–6, n=58), early–mid (LH+7–8, n=290), mid (LH+9–10, n=250), and late (LH+11–14, n=66) luteal phases. Expression of 13 established endometrial marker genes was measured using RTq-PCR together with two predefined gene-expression ratios associated with endometrial maturation.

#### Drift Changes Over the Luteal Phase

GOTFlow recovered coherent directional transitions across the luteal timeline, where transitions are inferred in a 8-dimensional learned latent embedding of the gene-expression profiles visualised in 2D using PLS (Fig. 3.A). These transitions appear as drift vectors describing transcriptional progression across states. Drift magnitude increased from early to mid-luteal phases, consistent with accelerated transcriptional reprogramming during decidual activation. Temporal markers including *CXCL14, GPX3*, and *DPP4* showed increasing drift contributions across states, whereas *DIO2* and *SLC15A2* contributed negative shifts consistent with their expected downregulation during progesterone-driven differentiation. Together these patterns indicate that GOTFlow captures transcriptional trajectories underlying the transition to a decidualised endometrial state.

**Figure 3.**
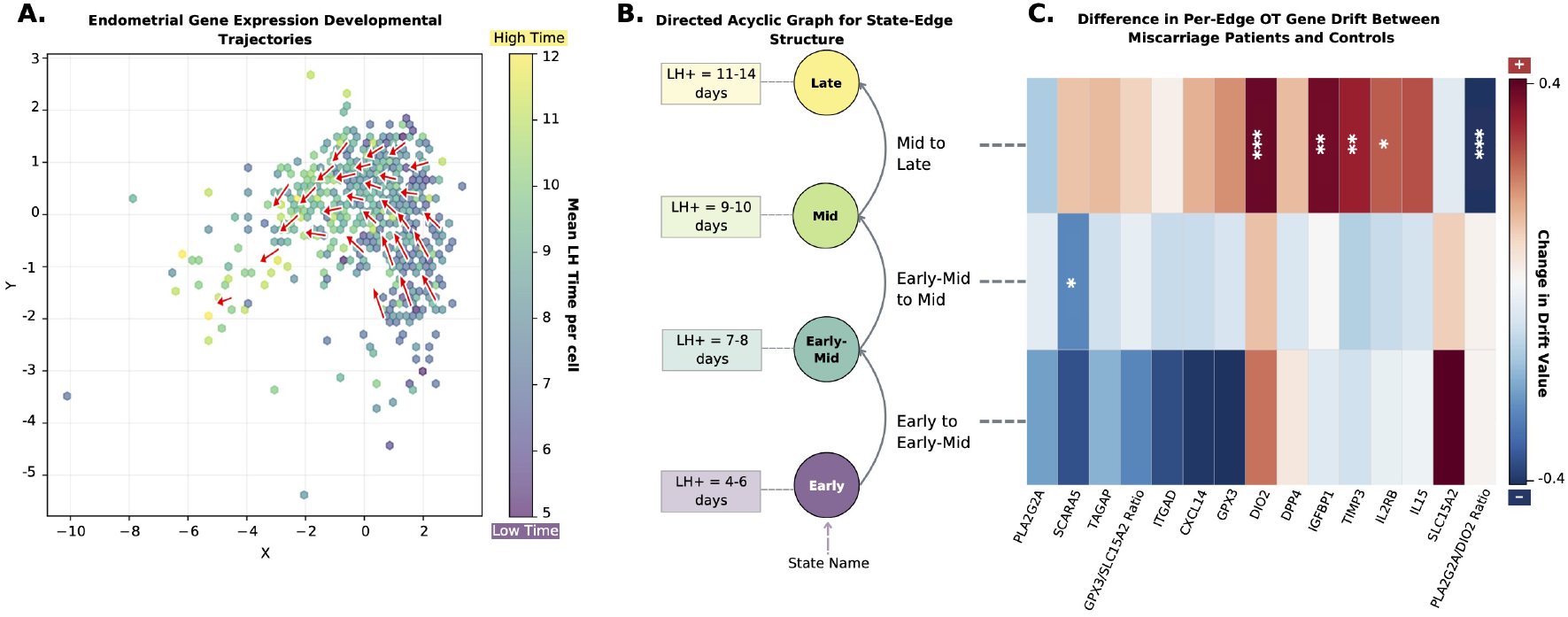
GOTFlow transport visualisation for the endometrial dataset. (A) Two-dimensional Partial Least Squares (PLS) projection of the learned GOTFlow latent embedding, coloured by mean target state; red arrows indicate inferred drift vectors between states. (B) Directed acyclic graph (DAG) specifying the state transitions used for modelling. (C) Per-edge feature-level drift differences between patients with a history of pregnancy loss and controls. Red indicates genes with increased expression shifts during transitions in miscarriage patients, whereas blue indicates decreased shifts. Statistical significance for difference in miscarriage patients is indicated by t-test p-values: * *p* < 0.05, ** *p* < 0.01, *** *p* < 0.001.

#### Reduced Drift Magnitude Characterises Miscarriage Patients

Miscarriage samples showed consistently lower drift magnitudes than controls, indicating slower progression through gene-expression state space during endometrial remodelling. This difference was most pronounced during the early–mid luteal transition (LH+7–8), when decidualisation initiates.

To quantify this effect, we fitted a Bayesian regression model:

Drift Magnitude *∼* Miscarriage History + LH + Age + BMI.

Miscarriage history was found to be significantly associated with reduced drift magnitude (*β* = −0.143 *±* 0.072, 90% CrI [−0.268, −0.031]), independent of age, BMI, and LH+ day (Table 1). This pattern is consistent with previous reports of delayed or impaired endometrial maturation in miscarriage-associated cycles (16). Architecture ablation analyses are reported in Supplementary Results (Table S2).

**Table 1.** Posterior summaries from the Bayesian regression model estimating drift magnitude, reporting posterior means, standard deviations, and 90% credible intervals (highest-density intervals) for all regression coefficients. The estimates indicate reduced drift in miscarriage samples compared with controls after adjustment for age, BMI, and LH+ day as credible intervals do not cross zero.

| | Mean ± std | 5% CrI | 95% CrI |
| --- | --- | --- | --- |
| Intercept | 2.183 ± 0.358 | 1.586 | 2.751 |
| Miscarriages | -0.143 ± 0.072 | -0.268 | -0.031 |
| LH | -0.038 ± 0.020 | -0.072 | -0.007 |
| Age | -0.011 ± 0.008 | -0.024 | 0.002 |
| BMI | 0.005 ± 0.007 | -0.016 | 0.006 |

#### Feature-Level Drivers of Impaired Decidualisation

Feature-level drift analysis, obtained by projecting transport barycentres back to gene space, revealed specific molecular programs underlying these altered trajectories (Fig. 3.C). The *PLA2G2A/DIO2* ratio, a marker of the receptive-to-decidualised transition previously linked to miscarriage (16), showed significantly reduced temporal change in miscarriage patients (FDR-corrected *p* < 0.05), particularly during later luteal stages. In contrast, *IGFBP1, TIMP3*, and *IL2RB* displayed increased drift contributions, consistent with dysregulated stromal responses during decidualisation. Together these findings indicate that GOTFlow identifies altered transcriptional trajectories associated with impaired endometrial maturation preceding reproductive failure.

### Modelling Risk Trajectories in Breast Cancer

Breast invasive carcinoma (BRCA) exhibits substantial heterogeneity in clinical outcome, motivating approaches that connect molecular variation to disease risk. We analysed bulk RNA-seq data from 1061 tumours in the TCGA-BRCA cohort (14), restricting the feature space to a curated panel of 608 prognostic genes from the Human Protein Atlas (20). Using these genes, we trained a survival risk model (2) to assign each patient a continuous risk score reflecting predicted survival. To define an ordered progression structure, the risk scores qunatiles were discretised into *K* = 5 states spanning low to high risk. We then applied GOTFlow to analyse molecular transitions underlying this risk continuum. It learned a 32-dimensional embedding and estimated OT couplings between adjacent risk states, with a 2D visualisation obtained via PLS projection. The resulting transport map (Fig. 4A) reveals coherent directional flow from lower-to higher-risk strata, indicating progressive molecular shifts accompanying increasing predicted risk.

**Figure 4.**
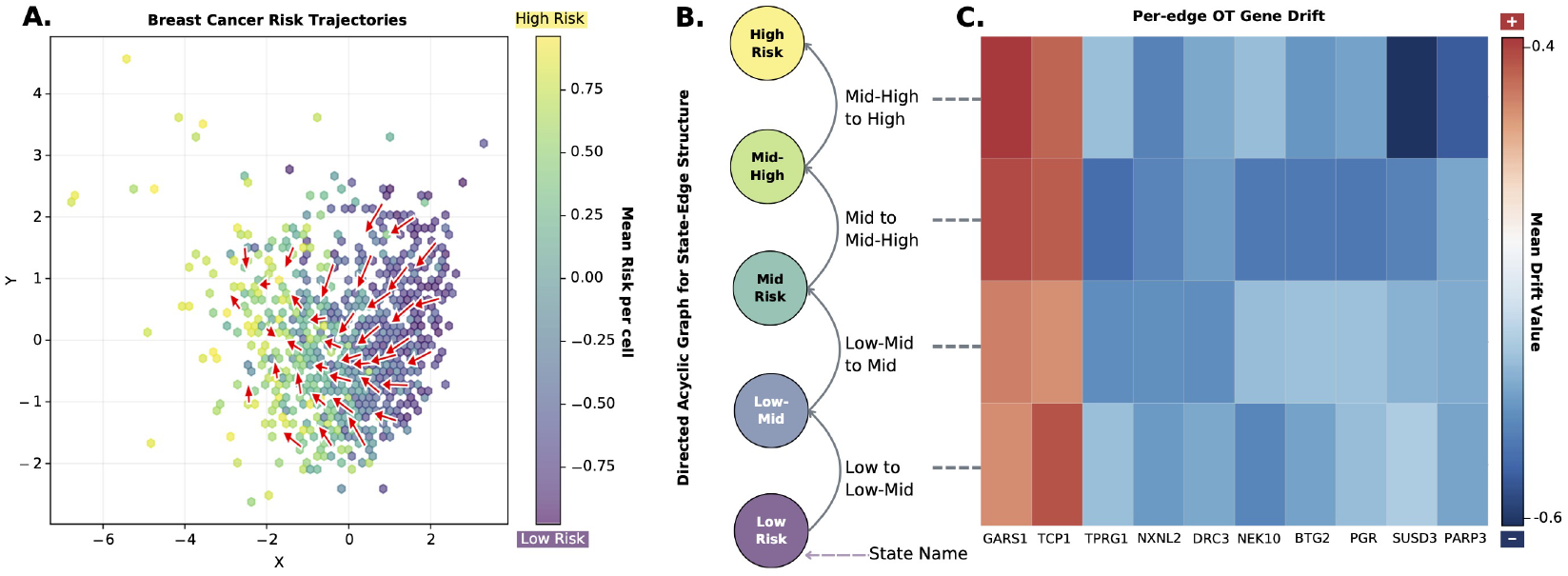
GOTFlow transport visualisation for the breast cancer dataset. (A) Samples are embedded in 2D using PLS and coloured by mean target per cell; red arrows indicate the inferred drift. (B) Directed Acyclic Graph (DAG) illustrating the state nodes and edges. (C) Per-edge feature-level drift heatmap showing OT-implied gene-expression shifts across transitions. Red indicates genes with increasing expression during transitions to higher-risk states, whereas blue indicates decreasing expression.

#### Barycentric Feature Drift Reveals Risk-Associated Gene Shifts

To identify genes driving these transitions, we analysed barycentric feature drifts derived from the OT couplings in the original gene-expression space. Fig. 4.C shows the ten genes with the largest transported-mass-weighted shifts across adjacent risk transitions (additional genes are shown in Supplementary Fig. S2). Clear directionality emerged: some genes showed progressively increasing expression along the risk trajectory, whereas others decreased consistently across states. Genes with increasing drift contributions included *TCP1* and *GARS1*, both previously linked to aggressive tumour biology. *TCP1* has been reported as recurrently altered in breast cancer and required for tumour cell growth (9), while *GARS1* promotes malignant phenotypes through proliferative signalling pathways (12). In contrast, genes with decreasing shifts included *SUSD3* and *NXNL2*, which have been associated with favourable prognosis or therapy responsiveness (15; 6).

Consistent with the OT-derived drift patterns, the median expression of these genes changed monotonically across ordered risk states (Supplementary Fig. S3). Genes with positive OT shifts increased as predicted risk increased, whereas genes with negative shifts decreased across states. A complementary univariate survival analysis of these OT-selected genes is provided in Supplementary Fig. S4, confirming that these features individually carry prognostic signal. Together these results indicate that GOTFlow recovers biologically meaningful transcriptional programs underlying the survival model’s risk stratification.

### Mouse Brain Transcriptional Dynamics in Prion Disease

Prion diseases are fatal neurodegenerative disorders characterised by progressive accumulation of misfolded prion protein (*PrP* ^*Sc*^) and widespread transcriptional disruption in the brain. To characterise the molecular trajectories underlying disease progression, we analysed a transcriptomic dataset of mouse brain samples collected across the incubation period following prion infection (11). In the original study, mice were inoculated intracerebrally with prion-infected brain homogenate, while genotype-matched controls received non-infectious homogenates and were sampled in parallel. Brain tissue was collected at 8–10 time points spanning asymptomatic to terminal clinical stages, resulting in a total of 450 Affymetrix Mouse Genome 430 2.0 microarrays. Expression data were processed using the original normalisation pipeline. To focus the analysis on genes exhibiting meaningful temporal variation, low-variance genes (variance *<* 0.5) were removed, leaving 99 transcriptional features for modelling. These features capture genes whose expression varies substantially over disease progression and therefore provide a compact representation of the underlying transcriptional dynamics.

#### Progression Trajectories Identify Disease-Associated Genes

To model transcriptional dynamics during disease progression, samples were grouped into five temporal bins representing successive stages of the incubation period. A directed state graph connecting adjacent stages was used to specify admissible transitions, and GOTFlow learned an 8-dimensional embedding, transport couplings and drift vectors between stages, revealing progressive shifts in gene-expression space. (Fig. 5A). Early transitions exhibit relatively small drift magnitudes, reflecting limited transcriptional perturbation during the asymptomatic phase, whereas larger drifts emerge at later stages as neuropathology develops. Near-zero drift between certain neighbouring states indicates periods of relative transcriptional stability, suggesting plateaus in the progression of molecular change. Together, these results delineate a temporal landscape of transcriptional perturbations associated with prion disease progression.

**Figure 5.**
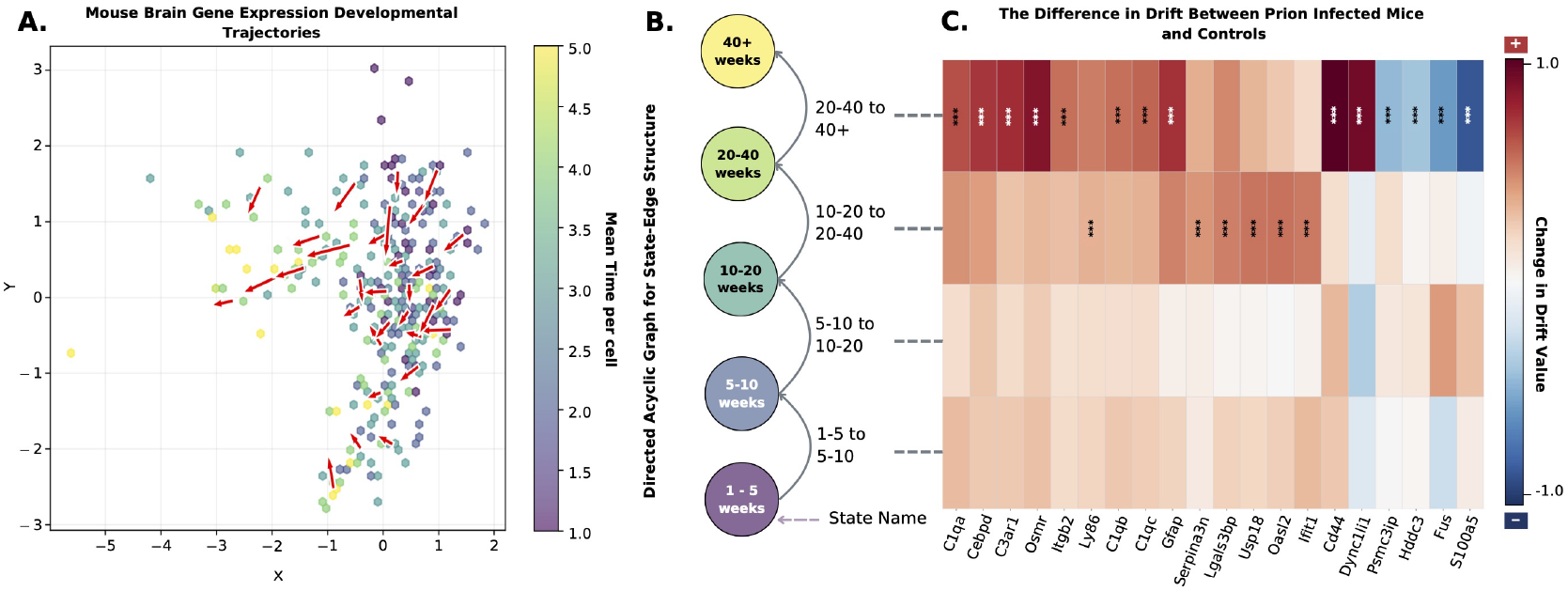
GOTFlow visualisation for the mouse brain dataset. (A) Samples are embedded in 2D using PLS and coloured by mean target per cell; red arrows indicate the inferred drift. (B) Directed Acyclic Graph (DAG) illustrating the state nodes and edges. (C) Per-edge feature-level drift (gene-shift) heatmap comparing prion-infected and control mice. Red indicates genes with increased OT-implied shifts when transitioning to the next state in prion-infected mice relative to controls, whereas blue indicates decreased shifts. Significant differences in infected patients are indicated with t-test p-values: *:p *<* 0.05, **:p *<* 0.01, ***:p *<* 0.001.

Feature-level drift analysis highlights genes that drive these transitions. By comparing inferred drift patterns between prion-infected and control mice, we identified gene-level disparities that pinpoint transcriptional programs specifically associated with prion pathology. Notably, *C1qa, Cd44*, and *Gfap* exhibited strong infection-specific directionality shifts (Fig. 5C). These genes are well-established markers of neuroinflammation and glial activation during prion disease, reflecting complement activation, astrocyte reactivity, and microglial responses to accumulating *PrP* ^*Sc*^ (19; 4; *3). The* concordance between GOTFlow-derived drift signatures and known prion-associated pathways demonstrates that the model recovers biologically meaningful transcriptional dynamics underlying neurodegenerative progression.

## Discussion, Limitations and Conclusions

GOTFlow provides a generalised framework for modelling directed transitions between biological populations using optimal transport. Rather than deriving trajectories from data, GOTFlow allows transitions to be specified through a user-defined directed state graph reflecting hypothesised biological relationships. Transport can be computed either in the original feature space or in a learned latent representation, while unbalanced optimal transport permits mass to appear or disappear between states, capturing changes in population composition. These properties make GOTFlow particularly suited to cross-sectional biomedical datasets and position it as a generalisation of optimal-transport-based analyses for graph-constrained modelling of population transitions. In synthetic experiments with known transition structure, GOTFlow recovered the correct directional dynamics, providing controlled validation of the learned embedding and transport objective. Applications to three biological domains (endometrial remodelling, breast cancer risk progression, and prion disease) further illustrate the promise of the framework by revealing biologically meaningful molecular shifts associated with reproductive failure, cancer progression, and neurodegeneration. Because this formulation differs from many trajectory or OT-based approaches that infer transitions or operate in fixed embeddings, direct benchmarking would require substantial methodological modification of those methods. The goal of this work is therefore not to claim superiority but to introduce and demonstrate the utility of this modelling framework.

Several limitations should be considered: As can be expected of any such an approach, inferred dynamics depend on user-defined state discretisation and transition structure, which encode assumptions about the underlying progression. Moreover, while GOTFlow highlights features associated with inferred trajectories, these relationships are observational and do not establish causality and experimental studies are required to validate the underlying mechanisms.

## Data availability

BRCA gene-expression data were obtained from TCGA via the Genomic Data Commons (https://portal.gdc.cancer.gov). Mouse brain data are available in ArrayExpress under accession E-MTAB-76. Endometrial data were provided by the Brosens Lab.

## Author contributions statement

F.M. conceived the approach, E.A. and G.W. designed and conducted the experiment(s), J.M. and J.B. collected the endometrial data. G.W. E.A. and F.M. wrote and all authors edited the manuscript.

## References

1. S Aldridge et al. Single cell transcriptomics comes of age. Nature communications, 2020.

2. E Alzaid et al. Automatic discovery of robust risk groups from limited survival data across biomedical modalities. Machine Learning with Applications, 2026.

3. GM Bentivenga et al. Diagnostic and prognostic value of plasma gfap in sporadic creutzfeldt–jakob disease in the clinical setting of rapidly progressive dementia. International Journal of Molecular Sciences, 2024.

4. BM Bradford et al. Cell adhesion molecule cd44 is dispensable for reactive astrocyte activation during prion disease. Scientific Reports, 2024.

5. C Bunne et al. Learning single-cell perturbation responses using neural optimal transport. Nature methods, 2023.

6. F Conte et al. In silico recognition of a prognostic signature in basal-like breast cancer patients. PLoS ONE, 2022.

7. Jean Feydy et al. Interpolating between optimal transport and mmd using sinkhorn divergences. In The 22nd International Conference on Artificial Intelligence and Statistics, 2019.

8. A Forrow et al. Lineageot is a unified framework for lineage tracing and trajectory inference. Nature communications, 2021.

9. S Guest et al. Two members of the tric chaperonin complex, cct2 and tcp1 are essential for survival of breast cancer cells and are linked to driving oncogenes. Experimental Cell Research, 2015.

10. W Hou et al. A statistical framework for differential pseudotime analysis with multiple single-cell rna-seq samples. Nature communications, 2023.

11. D Hwang et al. A systems approach to prion disease. Molecular systems biology, 2009.

12. Ealia Khosh K et al. Glycyl-trna synthetase (gars) expression is associated with prostate cancer progression and its inhibition decreases migration, and invasion in vitro. International Journal of Molecular Sciences, 2023.

13. G La Manno et al. Rna velocity of single cells. Nature, 2018.

14. J Liu et al. An integrated TCGA pan-cancer clinical data resource to drive high-quality survival outcome analytics. Cell, 2018.

15. I. Moy et al. Sushi domain containing 3 (susd3) and its role in breast cancer cell morphology, migration, and adhesion. Fertility and Sterility, 2011.

16. J Muter et al. Stalling of the endometrial decidual reaction determines the recurrence risk of miscarriage. Science Advances, 2025.

17. W Saelens et al. A comparison of single-cell trajectory inference methods. Nature biotechnology, 2019.

18. G Schiebinger et al. Optimal-transport analysis of single-cell gene expression identifies developmental trajectories in reprogramming. Cell, 2019.

19. RB Sim et al. C1q binding and complement activation by prions and amyloids. Immunobiology, 2007.

20. M Uhlen et al. A genome-wide transcriptomic analysis of protein-coding genes in human blood cells. Science, 2019.

