## supplementary for "GOTFlow: Learning Directed Population Transitions from Cross-Sectional Biomedical Data with Optimal Transport"

### Supplementary Results

### Synthetic Data Study

A synthetic dataset was generated to model a simplified differentiation trajectory with a single branching event. The simulation comprised 100 cells embedded in a 64-dimensional feature space and evolved over 150 discrete time points. A branching point was introduced at $t = 50$, dividing the process into a shared trunk phase and two post-branching lineages.

During the trunk phase, each cell's state vector $\mathbf{x}_t$ was updated iteratively as

$$\mathbf{x}_t = \mathbf{x}_{t-1} + \mathbf{d}_{\text{trunk}} + \boldsymbol{\epsilon}_t, \boldsymbol{\epsilon}_t \sim \mathcal{N}(0, \sigma^2 I),$$

where $\mathbf{d}_{\text{trunk}}$ contained three up-regulated and three down-regulated components.

At the branch point, cells were evenly divided between two descendant lineages. Both lineages inherited the terminal trunk states but diverged according to modified direction vectors, $\mathbf{d}_A$ and $\mathbf{d}_B$, constructed by adding three positive or negative directional components, respectively, to the trunk vector.

Feature analysis reflected the intended trajectory design: features 1–3 exhibited a consistent positive drift, and features 4–6 showed a decrease across the trunk phase. Following the branch, the additional perturbed features displayed the expected directional changes, confirming that the model correctly captured and interpreted lineage-specific trends (Supplementary Figure S1).

### Miscarriage Ablation Study

We conducted an ablation study to assess how feature maps and transport couplings affect estimated effect of miscarriage on drift across blur levels. Smaller blur values (controlled by epsilon) produce sharper transport plans that couple nearby samples in the latent space, whereas larger blur values yield smoother couplings that capture broader population-level structure. In the context of GOTFlow, this parameter therefore controls the scale at which transitions between state populations are resolved in the learned latent embedding.

Table S2 evaluates the effect of the Sinkhorn blur parameter and model components on drift estimation. The full model combining a learned representation with optimal transport produces the strongest and most stable drift estimates across blur values. Using the identity mapping weakens the effect, while replacing optimal transport with uniform couplings removes the signal entirely, indicating that the detected drift depends on structured transport alignment rather than simple averaging. As blur increases, drift estimates shrink toward zero due to the increasingly diffuse transport plans.

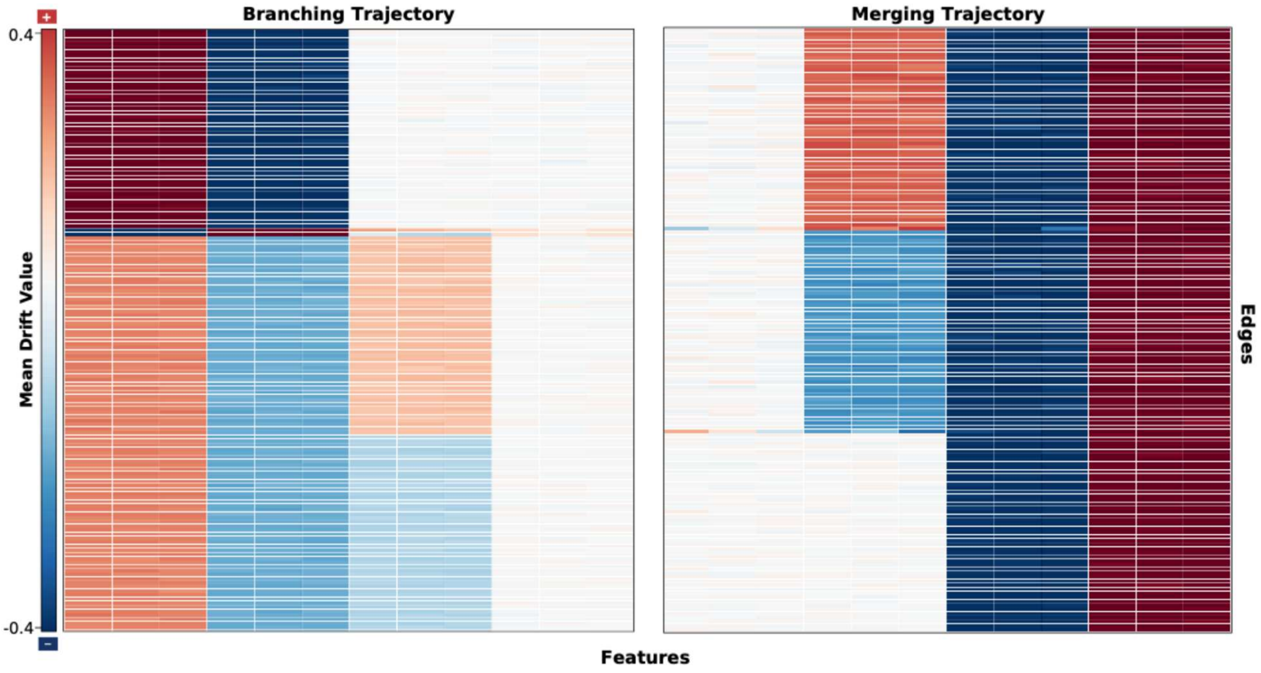


**Figure S1**: Per-edge feature-level drift heatmap for the synthetic dataset, for both branching and merging trajectories. Red indicates genes with positive OT-implied shifts (increasing expression) when transitioning to the next state, whereas blue indicates negative shifts (decreasing expression).

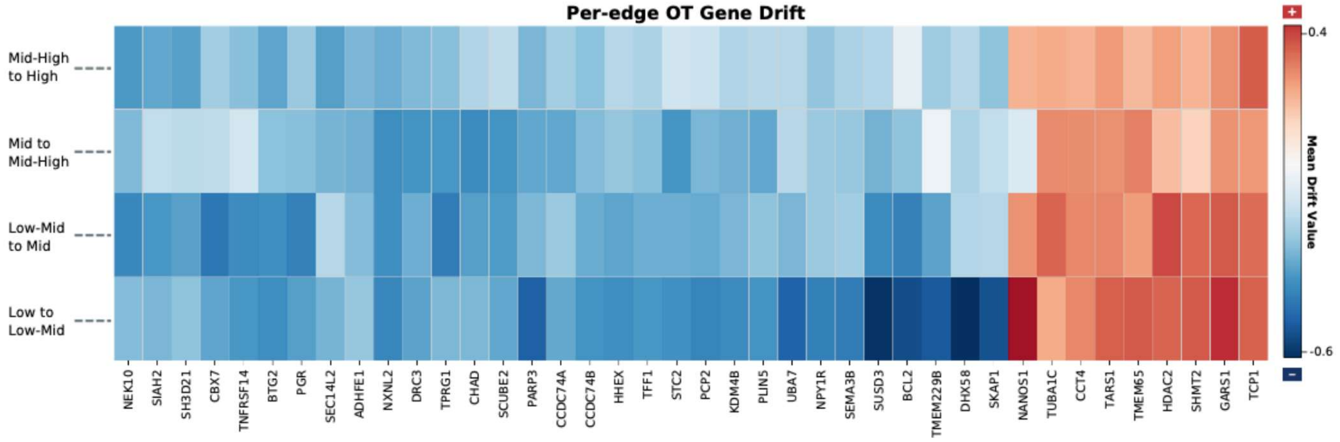


**Figure S2:** Per-edge feature-level drift (gene-shift) heatmap for the breast cancer dataset. Red indicates genes with positive OT-implied shifts (increasing expression) when transitioning to the next state, whereas blue indicates negative shifts (decreasing expression).

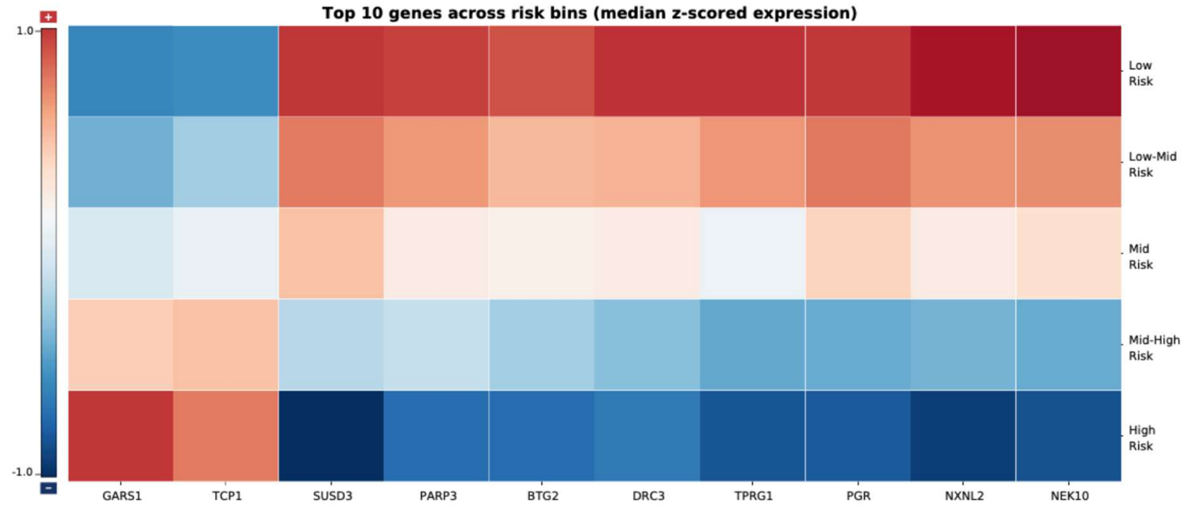


**Figure S3:** Expression patterns of top transport genes in the breast cancer dataset across ordered states. Heatmap showing the expression of the highest-transport genes across ordered states, summarised per state by median expression. The observed trends align with the OT-implied directionality, with genes exhibiting positive shifts increasing across states and genes with negative shifts decreasing.

### High-Transport Genes Show Progressive Expression Changes Across Risk States

We performed a univariate survival analysis to test whether the OT-selected genes were individually associated with patient outcome. This step provided an orthogonal check that the genes highlighted by GOTFlow were not only changing along the inferred trajectory, but also carried prognostic signal when considered one at a time. Using a Cox proportional hazards model fitted separately for each gene, we estimated hazard ratios (HRs) alongside statistical significance with multiple-testing control. A hazard ratio greater than one indicates that higher gene expression is associated with increased hazard (poorer survival), whereas a hazard ratio less than one indicates a protective association. We report the univariate Cox results as a forest plot in Fig.S4, showing the estimated hazard ratios with their confidence intervals for each gene. Consistent with the OT directionality, *TCP1* and *GARS1* exhibited elevated hazard ratios, while genes decreasing along the trajectory showed hazard ratios below one. All tested genes remained significant after false discovery rate correction (FDR-adjusted $p < 0.001$).

At the state level, we further evaluated whether adjacent risk transitions corresponded to measurable survival differences. Kaplan--Meier curves stratified by samples within neighbouring states (Fig.S5) showed significant separation for edges (2,3) and (3,4) ($p < 0.05$), indicating that later transitions captured clinically meaningful risk differences. Finally, to illustrate gene-level effects, we stratified patients into high- and low-expression groups using the cohort median for each gene and plotted Kaplan-Meier curves (Fig.S6). Separation between groups was assessed by a log-rank test, and all reported comparisons were significant ($p < 0.001$).

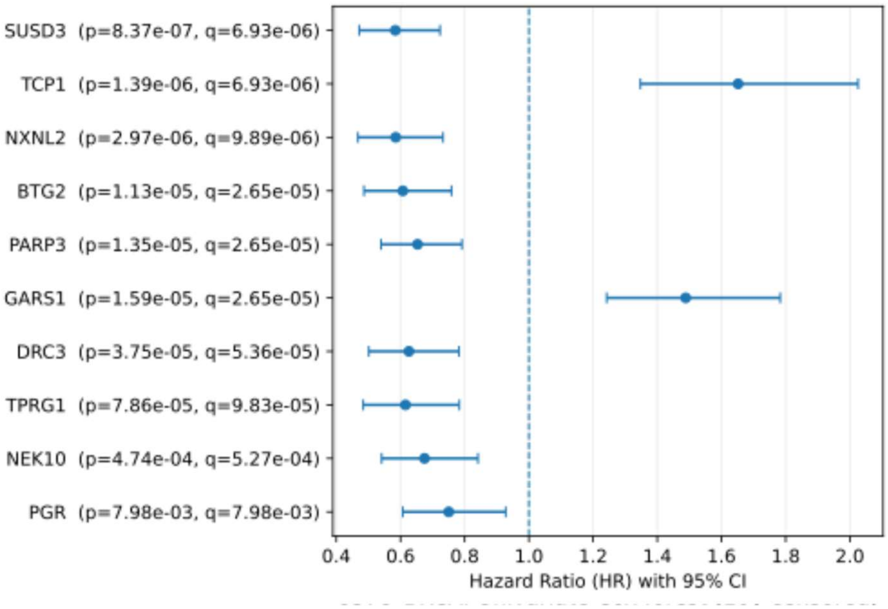


**Figure S4:** Univariate Cox Forest plot for high-transport genes in the breast cancer dataset. Forest plot showing hazard ratios (HR) with 95% confidence intervals from univariate Cox proportional hazards models fitted separately for each selected gene. $HR > 1$ indicates that higher expression is associated with increased hazard (poorer survival), whereas $HR < 1$ indicates a protective association. P-values were adjusted for multiple testing using false discovery rate (FDR)

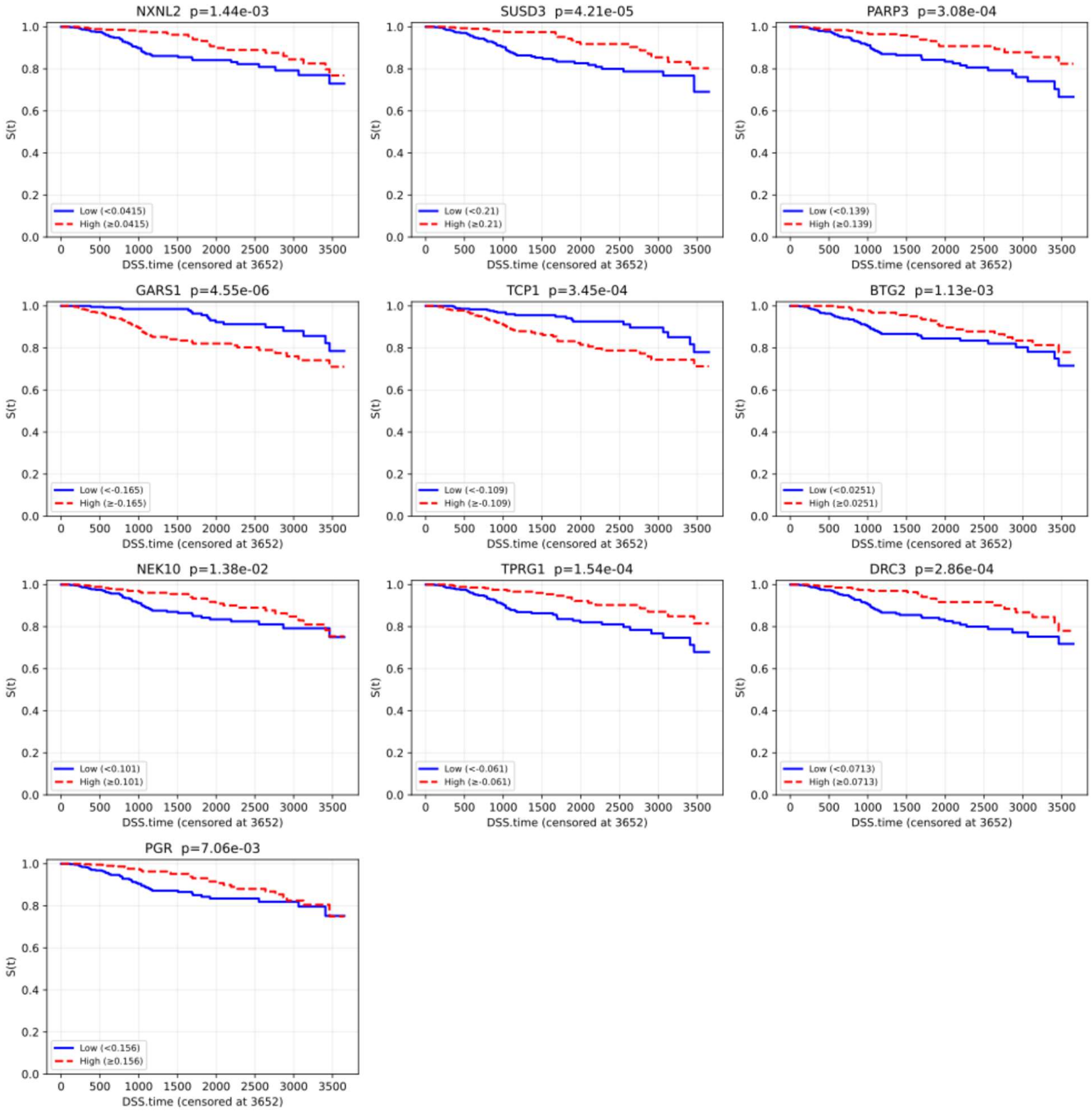


**Figure S4:** Kaplan-Meier curves for top genes in the breast cancer dataset stratified by median expression. For each gene, patients were divided into high- and low-expression groups using the cohort median expression as the threshold. Significance of separation was assessed using a log-rank test; all reported comparisons were significant ($p < 0.001$).

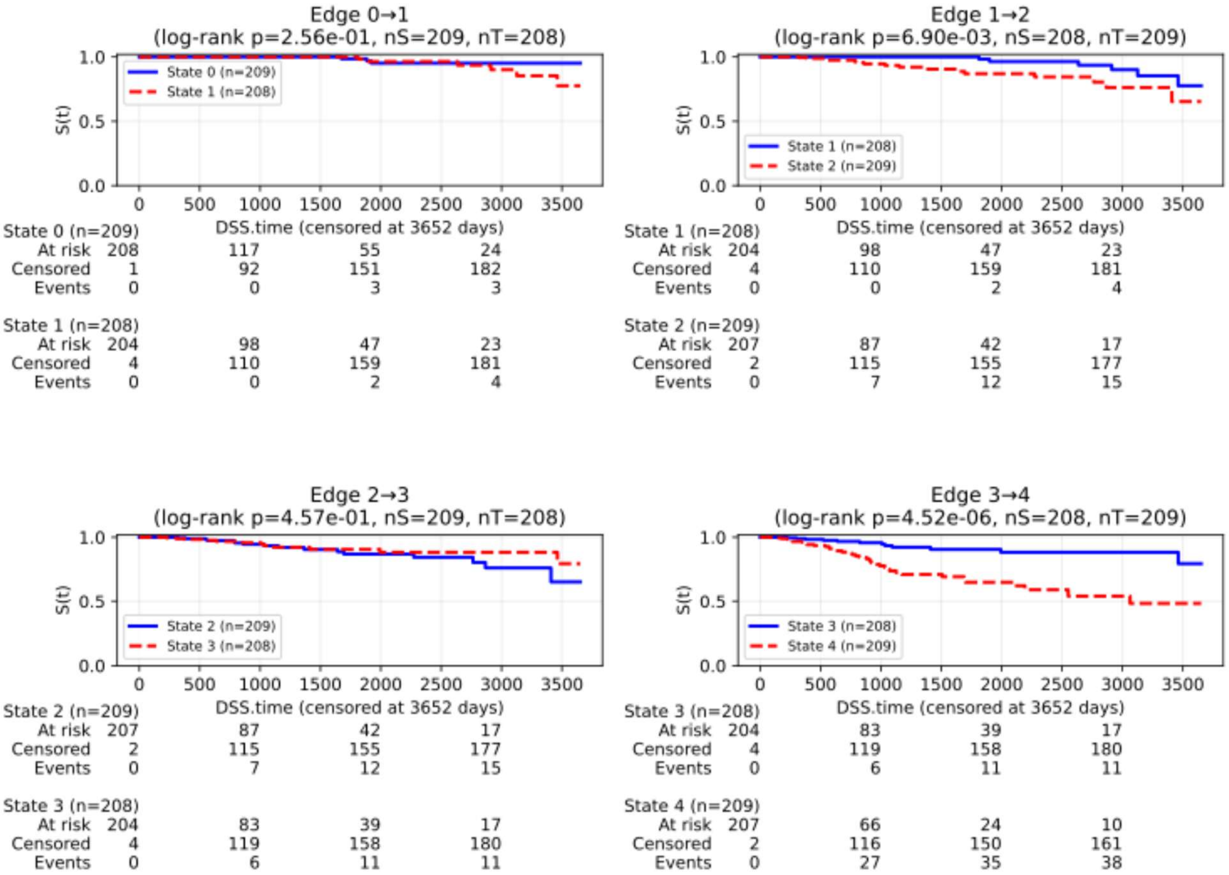


**Figure S5:** Kaplan-Meier curves to illustrate the survival differences between samples within adjacent edges for the TCGA-BRCA dataset. We note statistical significance ($p < 0.05$) for edges (2,3) and (3,4).

**Table S1.** Ablation study of GOTFlow architectures over synthetic data. Mean cosine similarities (± s.d.) between learned per-sample drift vectors and synthetic ground truth across model configurations.

| | Mean Cosine Similarity |
|---|---|
| GOTFlow | 0.720±0.152 |
| Without contrastive learning | 0.710±0.155 |
| With Balanced OT | 0.691±0.148 |
| Without Learned Embeddings | 0.364±0.887 |
| With Uniform OT | 0.365±0.887 |

**Table S2:** Ablation study of univariate Bayesian regression models for endometrial data across increasing blur levels, comparing learned versus identity feature mappings (Phi) and optimal transport (OT) versus uniform transport couplings, with posterior mean drift estimates and 90% credible intervals reported for each configuration.

| | Learned Phi and OT | | | Identity Phi and OT | | | Identity Phi and Uniform OT | | |
|---|---|---|---|---|---|---|---|---|---|
| Blur | Mean | 5% CrI | 95% CrI | Mean | 5% CrI | 95% CrI | Mean | 5% CrI | 95% CrI |
| 0.1 | -0.132 ± 0.065 | -0.237 | -0.026 | -0.126 ± 0.061 | -0.227 | -0.026 | -0.060 ± 0.123 | -0.262 | 0.139 |
| 0.2 | -0.15 ± 0.069 | -0.263 | -0.038 | -0.114 ± 0.053 | -0.201 | -0.027 | N/A | N/A | N/A |
| 0.3 | -0.187 ± 0.069 | -0.301 | -0.074 | -0.102 ± 0.057 | -0.194 | -0.008 | N/A | N/A | N/A |
| 0.4 | -0.142 ± 0.061 | -0.242 | -0.044 | -0.102 ± 0.058 | -0.199 | -0.008 | N/A | N/A | N/A |
| 0.5 | -0.143 ± 0.063 | -0.246 | -0.040 | -0.058 ± 0.058 | -0.154 | 0.037 | N/A | N/A | N/A |
| 0.6 | -0.084 ± 0.049 | -0.162 | -0.002 | -0.059 ± 0.068 | -0.171 | 0.053 | N/A | N/A | N/A |
| 0.7 | -0.106 ± 0.056 | -0.199 | -0.014 | -0.099 ± 0.073 | -0.219 | 0.022 | N/A | N/A | N/A |
| 0.8 | -0.07 ± 0.050 | -0.152 | 0.011 | -0.107 ± 0.077 | -0.234 | 0.018 | N/A | N/A | N/A |
| 0.9 | -0.042 ± 0.068 | -0.155 | 0.071 | -0.073 ± 0.070 | -0.188 | 0.041 | N/A | N/A | N/A |

### Policy for Acceptable Use of Large Language Models

We declare the use of LLM in the preparation of this study and manuscript under the acceptable uses, including:

- As an aid to correct written text (spell checkers, grammar checkers)
- As an algorithmic technique for research study
- As an evaluation technique (to assist in finding inconsistencies or other anomalies)
- Assist in code writing, and the code has been checked for correctness.
- Create documentation for code, and the documentation has been checked for correctness.
